# A Data Structure for real-time Aggregation Queries of Big Brain Networks

**DOI:** 10.1101/346338

**Authors:** Florian Ganglberger, Joanna Kaczanowska, Wulf Haubensak, Katja Bühler

## Abstract

Recent advances in neuro-imaging allowed big brain-initiatives and consortia to create vast resources of brain data that can be mined by researchers for their individual projects. Exploring the relationship between genes, brain circuitry, and behavior is one of key elements of neuroscience research. This requires fusion of spatial connectivity data at varying scales, such as whole brain correlated gene expression, structural and functional connectivity. With ever-increasing resolution, those exceed the past state-of-the art in several orders of magnitude in size and complexity. Current analytical workflows in neuroscience involve time-consuming manual aggregation of the data and only sparsely incorporate spatial context to operate continuously on multiple scales. Incorporating techniques for handling big connectivity data is therefore a necessity.

We propose a data structure to explore heterogeneous neurobiological connectivity data for integrated visual analytics workflows. *Aggregation Queries*, i.e. the aggregated connectivity from, to or between brain areas allow experts the comparison of multimodal networks residing at different scales, or levels of hierarchically organized anatomical atlases. Executed on-demand on volumetric gene expression and connectivity data, they enable an interactive dissection of networks, with billions of edges, in real-time, and based on their spatial context. The data structure is optimized to be accessed directly from the hard disk, since connectivity of large-scale networks typically exceed the memory size of current consumer level PCs. This allows experts to embed and explore their own experimental data in the framework of public data resources without large-scale infrastructure.

Our novel data structure outperforms state-of-the-art graph engines in retrieving connectivity of local brain areas experimentally. We demonstrate the application of our approach for neuroscience by analyzing fear-related functional neuroanatomy in mice. Further, we show its versatility by comparing multimodal brain networks linked to autism. Importantly, we achieve cross-species congruence in retrieving human psychiatric traits networks, which facilitates selection of neural substrates to be further studied in mouse models.

## Introduction

Recent brain initiatives, such as the Allen Institute (Oh et al. 2014; Hawrylycz et al. 2012; Lein et al. 2007), the Human Brain Project (Markram et al. 2011), the WU-Minn Human Connectome Project (Van Essen et al. 2013), and the China Brain Project (Poo et al. 2016), have accumulated large sets of brain data for neuroscience research. Visual analytics emerges as a promising tool to mine this multimodal neurobiological data for insight into the functional organization of the brain. Such technologies allow to directly explore the relation between genes, neuronal circuitry and brain function and can quickly add context to experimental findings. However, the major challenges for visual analytic workflows arise from accessing, fusing and visualizing spatial brain data, such as brain gene expression, structural and functional connectivity, and non-spatial data, like gene associated with a given brain function. A particular challenge when exploring such heterogeneous neurobiological data is the alignment of their spatial reference. Depending on the data acquisition technique, it can be volumetric or region-wise, and different resources are not necessarily in the same reference space. This can lead to time consuming workflows that involve manual aggregation of the data that do not work continuously on different scales.

The entry point for many neuroscience workflows are local brain regions/areas and/or gene expression sites that are linked to a specific brain function. Such sites typically are the results from neuronal recording, imaging, optogenetics and behavioral neurogenetic studies (e.g. amygdala subnuclei in emotional processing (Haubensak et al. 2010; Kim et al. 2017). However, there is a lack of tools to link these local region/area- and/or primary expression site-entry points into local, mesoscale and global genetic, structural and functional spatial network context. Such context ranges from local circuit and genetic data available as sets of primary gene expression sites (sites where the gene creates products, such as proteins; e.g. from databases (Lein et al. 2007) or user generated local transcriptomic data), mesoscale structural connectivity (regions where primary gene expression sites project to; e.g. from connectivity databases (Oh et al. 2014) or from user generated spatially registered viral tracing or CLARITY methods) to global functional networks (e.g. user generated immediate early gene activity tagging or fMRI). Thus, from the neuroscientist’s perspective, methods are needed to embed novel with existing data across local-global scales and heterogenous context. Bridging this gap is expected to create significant synergies for updating, mining, communicating and sharing brain data.

Spatial networks are organized as nodes and edges embedded in a space and equipped with a metric (Barthelemy 2010). In neuroscience, nodes represent regions/areas in the brain, while edges describe their structural, functional (Betzel and Bassett 2017) or genetic (Richiardi and Altmann 2015) relation. The size and complexity of these networks, with up to billions of edges, created a need for sophisticated data handling techniques to allow further analysis and exploration (Bassett and Sporns 2017).

Several interactive frameworks for querying connectomic data in neuroscience have been published in recent years. Allen Brain Institutes’s BrainExplorer as well as its web interface (Oh et al. 2014) can identify pre-computed incoming and outgoing connections of pre-defined locations (injection sites) and anatomical regions in mouse. Meso-scale source/target sites are visualized in 3D at voxel level and for quantitative examination ordered by brain region in a list. Although this is an easy-to-use tool for neurobiologists, results cannot be compared directly to other connectivity data or examined with respect to user-generated data. Other tools allow to locally explore the connectomes built by neurons traced on a single EM stack (volumetric electron microscopy) like CATMAID (Saalfeld et al. 2009) and ConnectomeExplorer (Beyer et al. 2013). They are working on a local level of a single network with a fixed scale. Similar accounts for Sherbondy et al. (Sherbondy et al. 2005), who used queries on volumes of interest and pre-computed pathways to explore diffusion tensor imaging data, and Tauheed et al. (Tauheed et al. 2013), who developed spatial management techniques for dense spatial neuron simulations.

Querying large-scale spatial networks is originally applied in different domains, particularly on transportation/road networks, but nevertheless underlies the same principles (Barthelemy 2010). Early approaches in optimizing local queries on road network data has proposed in 1997 by Shekhar and Liu (Shekhar and Liu 1997). In principle, they store network nodes, and respectively their edges, as adjacency list. The list is ordered by a space filling curve, so nodes that are spatially close are stored on the same disk page. This reduces I/O costs and therefore increases query speed. The data structure was further improved by Papadias et al. (Papadias et al. 2003) and Demir (Demir and Aykanat 2010) with a tree-like hierarchical structure to efficiently process range queries (perform queries in circular range around a query point) and successor retrieval operations (get all successors of a network node). The hierarchical structure refers herby only to the spatial domain-, and is otherwise not related to the network data.

Further techniques to speed up network queries can be found in the more general domain of graph computation (Pienta et al. 2015). A common method is the use of advanced caching/paging strategies to hold often accessed parts of a graph in memory (Kyrola, Blelloch, and Guestrin 2012; Han et al. 2013; Roy, Mihailovic, and Zwaenepoel 2013; Chi et al. 2016; Leskovec and Sosič 2016). Other approaches using memory mapping of large-scale graphs as edge list files to handle them on the disk programmatically as if they were in the main memory (Lin et al. 2014). This allows graph processing with billions of edges on consumer level computers and mobile devices (Lin et al. 2014; Lin, Chau, and Kang 2013; Chen et al. 2015). LLAMA (Macko et al. 2015) further uses compressed row storage to harness sparsity. Recent graph computing frameworks such as FlashGraph (Zheng et al. 2015) further facilitate solid-state disks in combination with minimization of I/O operations to perform out-of-memory graph analysis algorithms (Ai et al. 2017).

Despite their universal applicability, the lack of spatial optimization results in inferior performance for *Aggregation Queries*, i.e. aggregated connectivity from, to or between a set of nodes on spatial networks (see *Section Performance Evaluation*). Some of these implementations are tailored to unweighted, binary graphs (Han et al. 2013; Lin et al. 2014; Chi et al. 2016) that are unsuitable to be generalized to perform aggregation queries on weighted, i.e. non-binary, connectivity data.

To our better knowledge, there is currently no tool which combines those state-of-the art techniques to allow interactive exploration of multimodal, multiresolution neurobiological connectivity data on a “big data” level.

We meet this demand by proposing a data structure for integration and real time querying of heterogeneous large-scale connectivity matrices at multi-scale voxel and region level by exploiting the hierarchical organization of brain parcellations in combination with spatial indexation.

Region-wise (e.g. resting state functional connectivity) or voxel resolution (structural connectivity, spatial gene expression correlation) connectivity data is aggregated hierarchically, to bridge the gap between different scales and resolutions. The hierarchies are anatomy-driven and can be flexibly generated for different ontologies and their related spatial region annotations. On the lowest level of these hierarchies, high resolution, voxel-wise connectivities with billions of edges (matrices with hundreds of gigabytes) are stored on hard disk in spatially organized indices for high-speed data access. Therefore, aggregated connectivity from, to or between brain areas can be retrieved, from voxel-level to large anatomical brain regions, in an instant.

For direct correlation of different connectivity data at voxel level, we are expecting data residing in the same spatial reference brain space, i.e. registered to a the same (multi-resolution) standard brain^1^. However, the dual indexing strategy allows us also to easily integrate and correlate data available only at region level with voxel wise data within the same brain space, but in principle also across brain spaces at region level if region correspondence is known. Data from public resources can be easily integrated in our data structure as well as private data generated during experiments in the lab.

We demonstrate the practical significance of this tool by presenting use cases reproducing recent biological findings by performing data integration and interactive queries on heterogeneous neurobiological data from mice and humans provided by large scale brain initiatives.

While traditional visualization tools have to be installed locally (M.-R. Xia, Wang, and He 2013) there is a trend towards developing web-based services. They have the potential to integrate publicly available and individually administrable data repositories to facilitate integrated workflows, complying with the shift of brain connectomics into the “big data” era (M. Xia and He 2017) and a complementation of experimental science by data driven science. However, current online interfaces often lack advanced visualization techniques (Brown and Van Horn 2016) or do not allow for interaction with the visualization (Sherif et al. 2015). A drawback of those solutions is that they do not allow to privately explore or analyze own data in context of the publicly available data repositories without disclosing the data. Therefore, we created a web-based, interactive local 3D segmentation on visualized data to define volumes of interest (VOI) that can be used to query on user-selected connectivity data sets accessible via our data structure. Result is the cumulative voxel-wise connectivity of the selected VOI that is visualized as intensity volume in the 3D rendering. This kind of interaction allows the researcher to relate integrated resources, for example incoming/outgoing connectivity on voxel-level, directly to spatial data like gene expressions.

In general, the proposed data structure allows handling data of different modalities delivering volumetric and/or connectivity data, which can be used for experimental hypothesis finding. The presented framework is applicable for multilevel functional predictions and extends its relevance across species. Therefore, it is suitable for virtual screening of complex networks, like those linked to psychiatric disorders, to functionally dissect the corresponding neural correlates in mouse.

## Materials and Methods

### Data

The data relevant for our system can be divided into three types, which in principle can stem from any species or modality:

*A hierarchical definition of brain regions and their associated positions on a reference brain*. This is basically a hierarchical parcellation of a given standard brain and its related ontology. A hierarchy is generally starting with the whole brain divided iteratively into sub-regions, where the lowest level contains the highest resolved regions. These regions have a dense voxel-level representation in the brain for mouse and a set of MNI152 space coordinates for human (representing biopsy sites). We exemplarily use the Allen Mouse Brain Atlas (AMBA) ontology with 1288 regions on 5 levels (Lein et al. 2007) on a 132×80×114 voxel space, and Allen Human Brain Atlas (AHBA) ontology with 1840 regions also on 5 levels (Hawrylycz et al. 2012) on 3600 MNI152 coordinate space (representing biopsy sites).

*Connectivity data* is given as weighted adjacency matrices. Rows/columns represent the connectivity strength between brain areas on different scales (voxel or region-wise). The weights can be in any range, positive or negative. In the context of the use-cases described in this paper, we used three different types:

1. Structural connectivity: In Ganglberger et al (Ganglberger et al. 2017) we compiled a voxel-wise structural connectivity matrix that shows the projections (efferent neurons) of ∼15% of the brain from *AMBA* and respectively how voxels in a 132×80×114 mouse brain (100 micron resolution, i.e. the side of a voxel has a length of 100-microns) are structurally connected (details in Supplementary Note 1). The 67500 × 450000 directed connectivity matrix is stored as an uncompressed 91.5 gigabyte *CSV* (comma separated value) files. Weights are normalized to range between 0 and 1.
2. Functional connectivity: Functional connectivity, representing correlation of BOLD fMRI signal shows the functional association of brain regions for specific tasks or resting state. We used a resting state connectome for human (Van Essen et al. 2013) and experimental mouse data, which is only region-wise available (∼80 regions). Weights are undirected and represent positive correlation coefficients between 0 and 1.
3. Spatial gene expression correlation networks: Correlated gene expression networks quantifies tissue-tissue relationships across genes (Lein et al. 2007; Richiardi and Altmann 2015). Detail on matrix creation can be found in Supplementary Note 1. The data consists of a 60000×60000 undirected connectivity matrices for mouse, that show the transcriptional similarity for a specific gene sets and 3600×3600 for human (Hawrylycz et al. 2012). The mouse data has a resolution of (67×41×58 mouse brain, 200-micron resolution), and is about 12 gigabyte as uncompressed *CSV* in size. The data consists of undirected weights, showing positive correlation coefficients between 0 and 1.

A *Volume of Interest* (*VOI*) is a spatially related set of coordinates in a reference space. These can be arbitrary selected voxels of a user or a brain region. A *VOI* defines an area in the brain, where a user is interested in the aggregated source or target connectivity of its individual points.

### Managing and Aggregating Hierarchical Connectivity Data

Querying huge graphs, with billions of edges is time consuming since connectivity data increases its size quadratically to the number of nodes and requires special solutions for fast access. This data, represented as connectivity matrices (weighted adjacency matrices) can easily grow up to hundreds of gigabytes, so keeping them in memory of standard PCs becomes infeasible.

When operating on different anatomical scales it is necessary to perform cumulative operations on the connectivity matrices. In this case large parts of the connectivity matrix needs to be loaded and aggregated (e.g. calculate region-wise connectivity from voxel-wise connectivity). Examples for aggregated connectivity, are the projection strength between brain regions, represented by the sum of their structural connectivity (Oh et al. 2014), or the average functional correlation between brain regions in task fMRI.

Furthermore, connectivities are not necessarily available on similar resolution. For example, in *Section Data*, we described structural connectivity on a 100-micron resolution, functional connectivity on a region level, and gene expression correlation on 200-microns. Up-sampling functional connectivity and gene expression correlation to 100-microns would require more storage space, while down-sampling to 200-micron or region-level loses information. Therefore, a mapping of the connectivity to a common standard brain space is necessary.

The data access structure we propose is tailored to take advantage of sparseness, anatomical or hierarchical parcellations, and spatial organization of the data, which, to our best knowledge, standard graph managing frameworks such as graph databases are not optimized for.

To allow interactive (real time) exploration of the brain connectivity space, the purpose of the data structure is retrieving aggregated source or target connectivity of specific *VOI*, such as anatomical regions or arbitrary user defined areas, on a voxel- or region-level in an instant. These *Aggregation Queries* are executed on connectivity matrices, which we define as weighted directed adjacency matrix

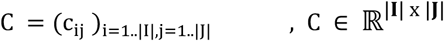

of a graph, where the rows **I** correspond to outgoing-, and the columns **J** to incoming edges of spatial regions, defining the spatial and/or anatomical resolution of the respective connectivity data (can be voxel level) in the discretized standard brain space **B** = {**p**_x_}_=1..n_, **p**_x_ ∈ ℝ^3^.

Here and in the following, the term *region* refers to a spatial related set of positions in a standard brain space such as anatomical brain regions, a group of voxels or a single voxel. Furthermore, we assume the standard brain space to represent the highest occurring resolution of all data to be queried.

### Spatial Mapping between Connectivity Matrices and Brain Space

We define the spatial association of the rows, respectively columns of the connectivity matrix **C**, to be a set of ordered disjunct sub-regions (i.e. from anatomical regions or voxel-level), so

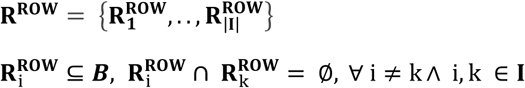

and respectively column associations **R**^**COL**^

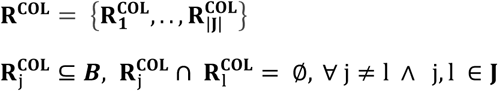

Note that **C** represents voxel-wise connectivity if

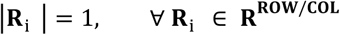

To directly associate spatial positions in brain space **p**_x_ ∈ **B** with rows and columns of **C**, i.e. incoming/outgoing connections we define the following mapping. At first, we map positions in brain space **p**_x_ to the indices of brain regions contained in **R**^**ROW**^ and **R**^**COL**^

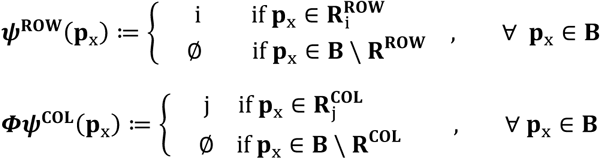

This creates non-unique mappings of arbitrary *VOI* in brain reference space **V** ⊆ **B** to rows/columns (i.e. a set of position in the brain space can point to multiple rows or column indices)

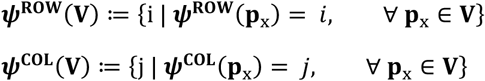

Please note that specific indices in the resulting set might be represented more than once, i.e. if there are m voxels in **V** laying in region **R**_i_, than i is m times present. This allows the usage of this mapping for aggregation of connectivity.

Vice versa, we map a set of indices to the union of their corresponding regions. As the voxel wise representation of regions in standard brain space is known, this generates a representation of connectivity at highest voxel resolution independent from the resolution or underlying parcellation of the original connectivity data.

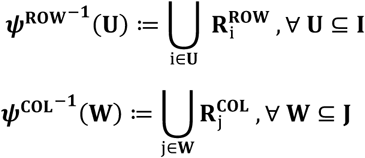

Therefore, a connection c_ij_ might represent equal connections of several points in brain. This has several advantages (see Figure 1):

**Figure 1.**
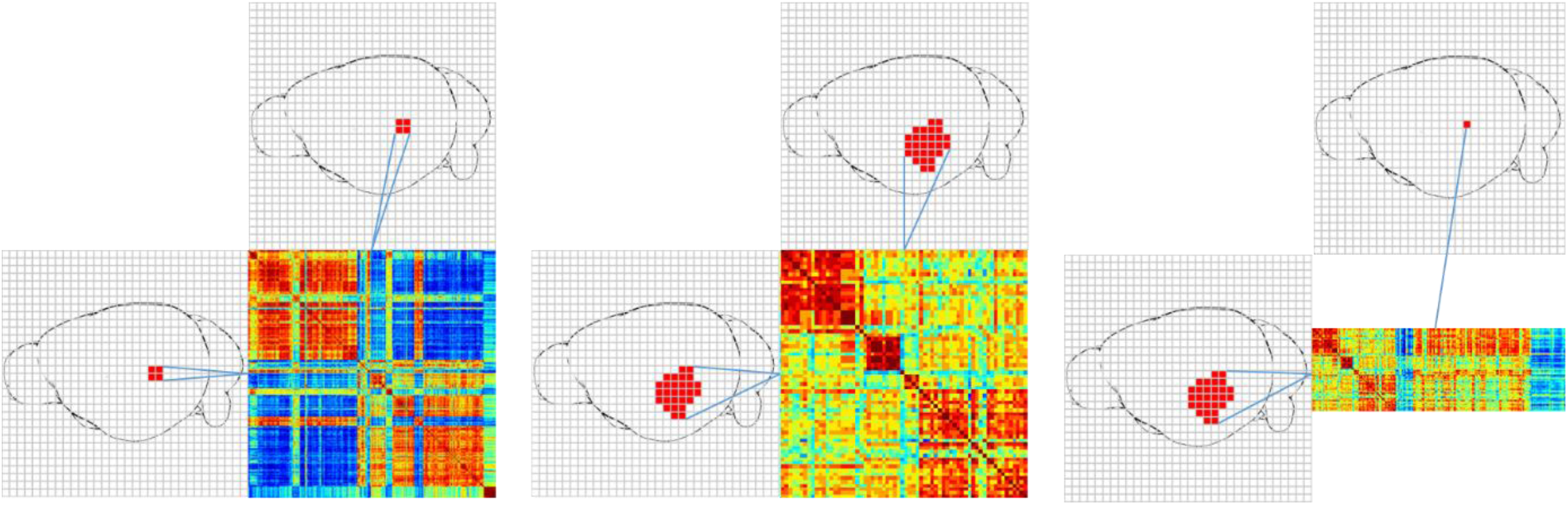
A: Connectivity matrix in a 4 times lower resolution than the reference brain space. Therefore every row/column is associated with 4 voxels. ***B***: Region-wise connectivity matrix. Every row/column is associated with voxels that form brain regions. ***C***: Region cache. Preprocessed aggregated outgoing connectivity for brain regions on voxel level.

1. Compare connectivity data defined on different resolutions of the standard brain: 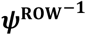 and 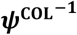 define the relation of rows respectively columns to voxel at a certain resolution. Since the overlap of regions with standard brain space is known, this enables a comparison of connectivity matrices in respect to different brain parcellations and/or different resolution. If the resolution is smaller than the reference space, this mapping would represent up-sampling (see Figure 1A).
2. Map region wise connectivity to voxel level: Nodes of a connectivity matrix can also represent (anatomical) brain regions to store region-wise connectivity data. Using 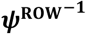 and 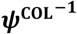 allow a retrieval of the data in voxel-wise brain space and therefore also the comparison of connectivity in respect to different brain parcellations (see Figure 1B).
3. Building caches: This technique can also be used to store precomputed data, such as connectivity of brain regions (from voxel level data) or pyramids representations with lower resolution (like an image pyramid). Although this increases the required storage, it improves scalability (see Figure 1C).

### A Dual Data Stucture Strategy for Aggregation Queries

*Aggregation Queries are defined as follows.* Let **V** ⊆ **B** be a *VOI*. The result of a target aggregation query is the cumulated outgoing connectivity for every position in space **B**

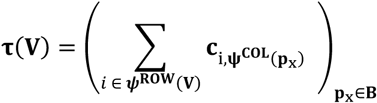

and the result of a source aggregation query the cumulated incoming connectivity for every row

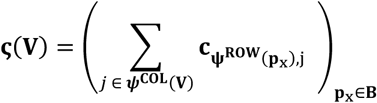

We are proposing a dual strategy unifying two complementary data structures to efficiently realize Aggregation Queries. The *Connectivity Storage* handles the data access for the *Aggregation Queries*, and the *Region-Wise Connectivity* in a Graph-Database manages queries on (anatomical brain-)region level. Figure 2 gives an overview of the overall system. Incorporating a connectivity matrix into our data structure begins with a preprocessing, that harnesses spatial-organization of the data (Figure 2 (1)) and uses row-compression to minimize disk-space (and therefore reading-time for queries) (Figure 2 (2)) to create a *Connectivity Storage File*. Region-wise connectivity of a hierarchical anatomical brain-region parcellation is precomputed and stored in a graph-database Figure 2 (3)). To further improving query performance, *Connectivity Cache Files* are created, that stores pre-computed connectivity for faster data access (Figure 2 (4)). Voxel-wise connectivity can then be queried from cache files and *Connectivity Storage Files* (Figure 2 (5)), region-wise connectivity form the graph-database (Figure 2 (6)). Preprocessing (Figure 2 (1,2,3)) is further described in the following subsections (*Connectivity Storage, Region-wise Connectivity Database*), and cache-creation as well as querying (Figure 2 (4,5,6)) in subsection *Implementation.*

**Figure 2:**
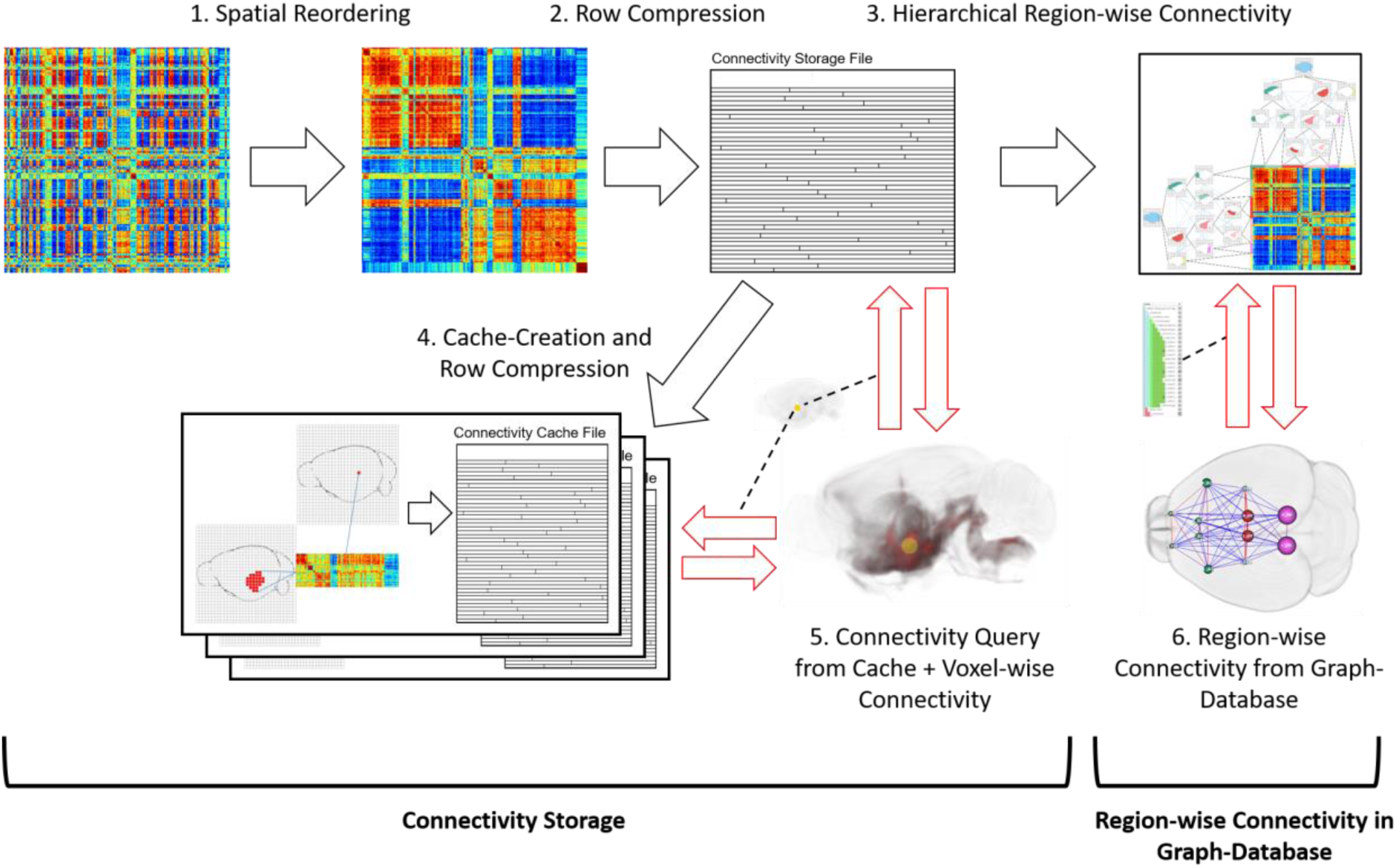
Overview of Connectivity Storage and the Region-wise Connectivity in the Graph-Database. Black arrows: Preprocessing of the data. **1.** Spatial Reordering of a (voxel-wise) connectivity matrix with a space filling curve. **2.** Row-wise compression of spatially-ordered connectivity matrix. 3. Gneration of hierarchical region-wise connectivity and storage in graph-database. **4.** Cache creation (preprocessed voxel-wise connectivity for predefined regions), and storage with row compression. Red arrows: **5.** Querying a *VOI* (yellow circle) on *Connectivity Cache Files*, then on *Connectivity Storage File*, resulting in aggregated connectivity (red). **6.** Querying connectivity between preselected brain regions (from a hierarchical parcellation), resulting in a region-wise connectivity graph.

#### Connectivity Storage

Since *Aggregation Queries* involve the reading and aggregating of whole rows or columns of connectivity matrices, we use a row-wise storage scheme. Allthough edge lists are popular for many graph management tools (Lin et al. 2014), which store connections in a <source node, target node, value> combination, they create a significant storage overhead for dense connectivity matrices.

Reducing data size allows higher query speed, since fewer data needs to be read. Therefore we apply a row-wise compression, that exploits potential sparseness of the data. First, the rows and columns of **C** are ordered by a space filling curve (Hilbert 1891) to preserve locality. The reordering causes sparse/dense areas to cluster within each row/column, since the connectivity of a region/voxel is not randomly distributed over the brain, but spatialy related. Then, a compressed row starts with the column index of the first non-zero value (*NZV*), the amount of *NZV* to follow, and the following *NZV*s. This is repeated similarly with the column index of the next *NZV* until the end of the row is reached. To identify each row in the file, an additional mapping

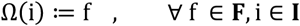

needs to be created, depicting the beginning of each row to their position f in the file **F**. A connection c_ij_ can be identified by going to the corresponding position of the i-th row f, and reading the j-th value from the row-wise compression. Figure 3 illustrates this process.

**Figure 3.**
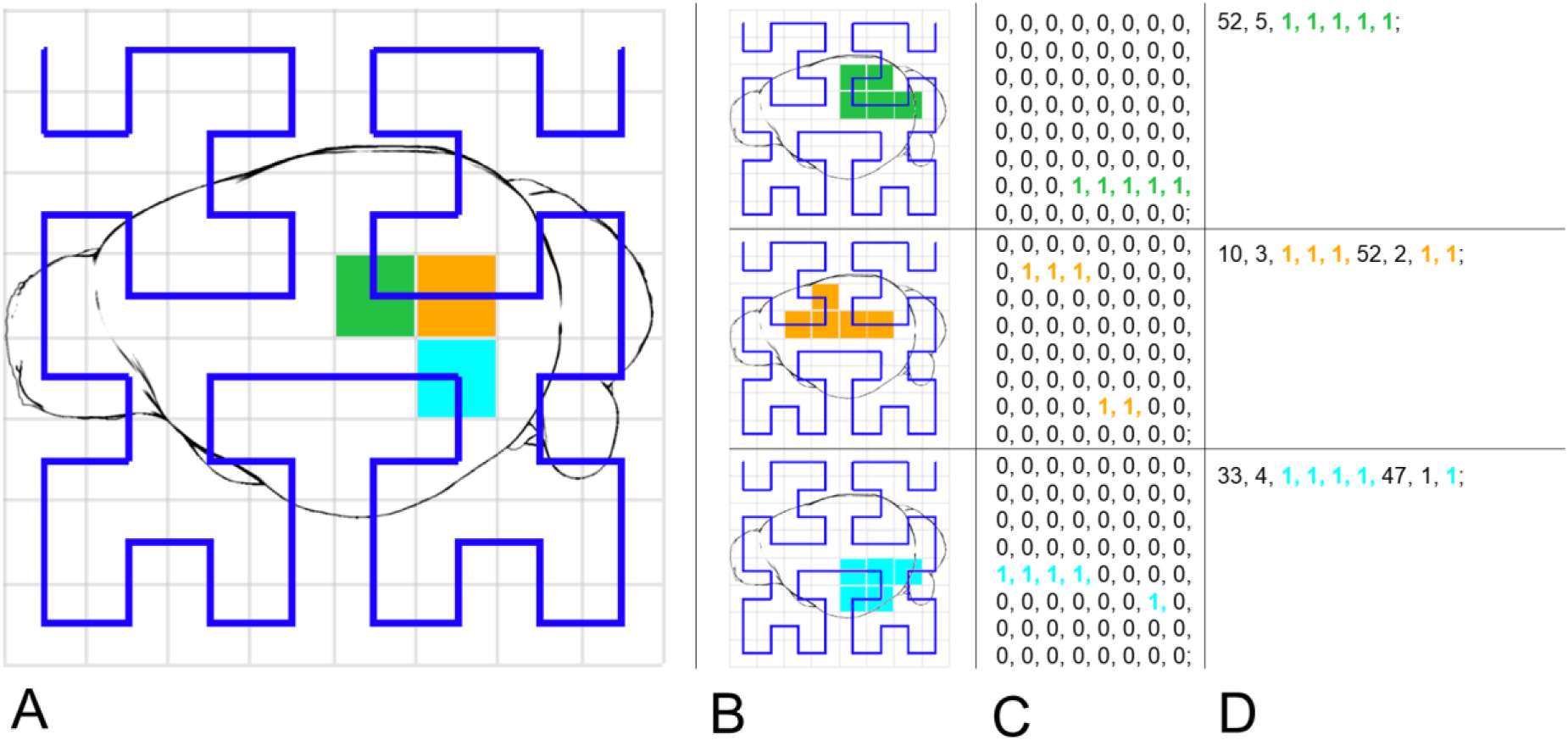
A: A mouse brain with overlayed Hilbert curve (blue), mapping the space to a one-dimensional space ***B***: The outgoing connectivity of the voxels corresponding to the colored voxels in *A.* Those would represent 3 rows in a connectivity matrix. ***C***: The connectivity of B along the hilbert curve (for simplicity in this example, connectivity is either 0 or 1). ***D:*** Row-wise compression of *C.* The compression can be read in this way: On the 52^nd^ position, 5 *NZVs* are following (green). On the 10^th^ position, 3, and on the 52th position 2 *NZVs* are following (orange). On the 33^rd^ position, 4 *NZVs* and on the 47^th^ position, there is 1 *NZV* following (cyan).

Other compression methods would also reduce the data size, but would not allow to directly access single rows without decompression of the whole file or significant parts of the file (Barrett et al. 1994).

For every connectivity matrix, we create a separate *Connectivity Storage File*, consisting three indizes as header (*FILE*: **Ω**(**i**)*, ROW:* ***Ψ***^**ROW**^(**V**), *COLUMN*: ***Ψ***^**COL**^(**V**) followed by compressed rows (Figure 4B). Even after compression, the compressed file does not necessarily fit into memory, especially when one wants to query on multiple connectivity matrices. We use memory mapping (*MMap*) to map the file into virtual address space. This allows programmatically access rows if they were in the main memory, without overhead of system calls. Further, the OS employs paging strategies, such as read-ahead paging. When performing a query for outgoing connections, the rows can be read in the order of their position in the file, and directly profit of read-ahead paging of the operating system to reach near-sequential reading speed. This additionally exploits the spatial organization of the data, that has been created with the ordering by space filling curve (see Figure 4A). Multiple connectivity matrices can then be queried sequentially without loading the whole matrices into memory. Note that a connectivity matrix of a directed graph needs an additional transposed *Connectivity Storage File* to query incoming connectivities (for undirected graphs, outgoing afnd incoming connections are equal due to symmetry).

**Figure 4.**
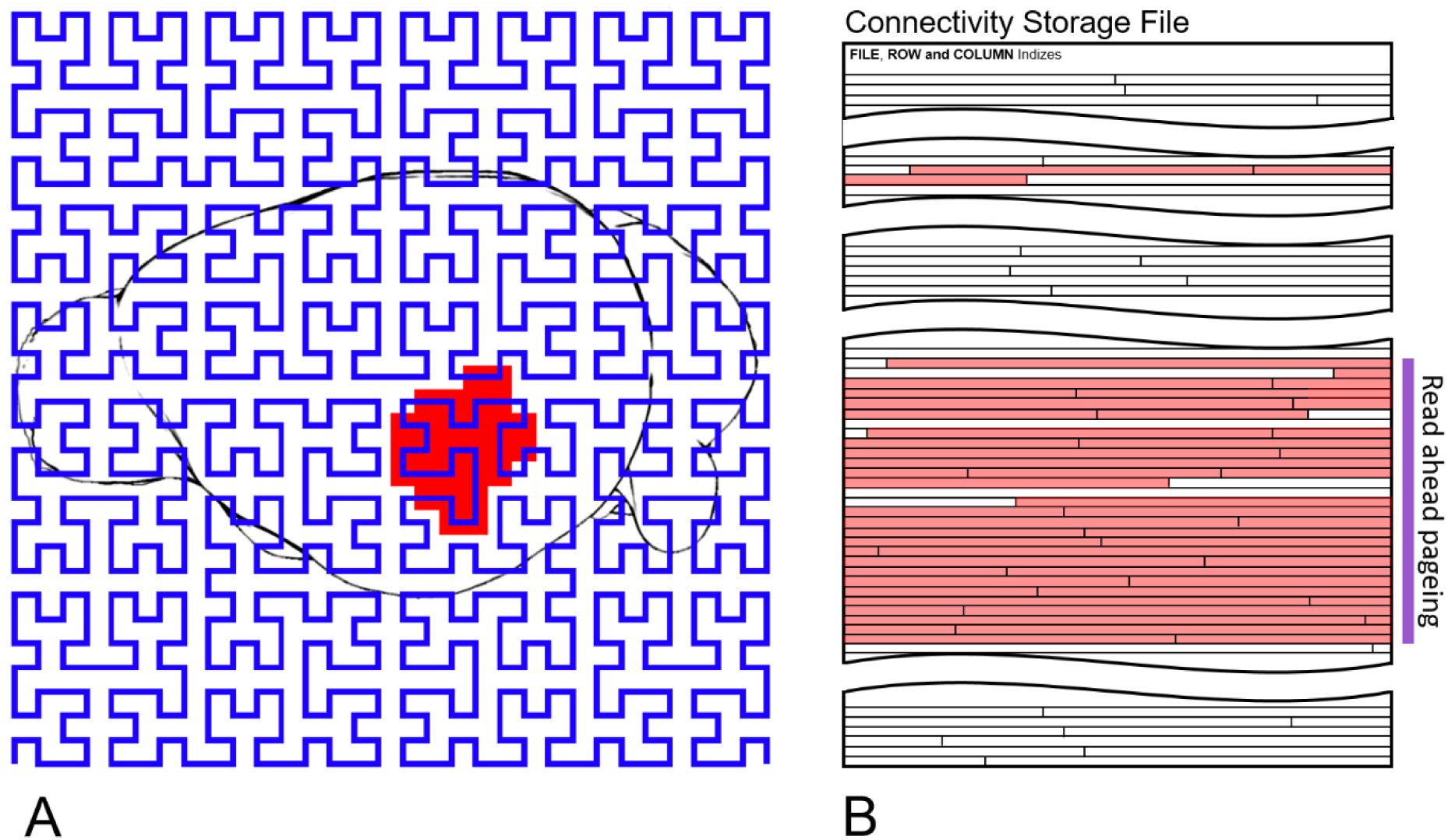
A: Brain Space overlayed with Hilbert curve (blue) and a *VOI* (red) that is queried for outgoing connectivity, ***B:*** Connectiviity matrix file (rows ordered by hilbert curve). Red-blocks represent rows that are read in order to get outgoing connectivity of *VOI* shown in *A*. Blocks can be read sequentially. Purple rows profit by read ahead paging.

#### Region-Wise Connectivity Database

On higher levels, the (anatomical brain-)region level, aggregated connectivity of a region consists of the connectivity of its subregions (and on the lowest level voxel-wise connectivity). When looking at brain wide region-wise graphs, it is not feasible to read the entire *Connectivity Storage* and compute the connectivity hierarchically at runtime. It would be to resource consuming for real-time computation. Instead, we compute it once when the *Connectivity Storage* is created, the resulting region-wise hierarchical connectivity is stored in a graph-database. The region-wise connectivity is computed recursively bottom up: First, the lowest level regions are aggregated from the *Connectivity Storage*, then the regions above are aggregated by their levels below until the top of the hierarchy. We further compute the connectivity between the levels in a similar way. Therefore, it is not necessary to compute any region-level connectivity at runtime (Figure 5).

**Figure 5:**
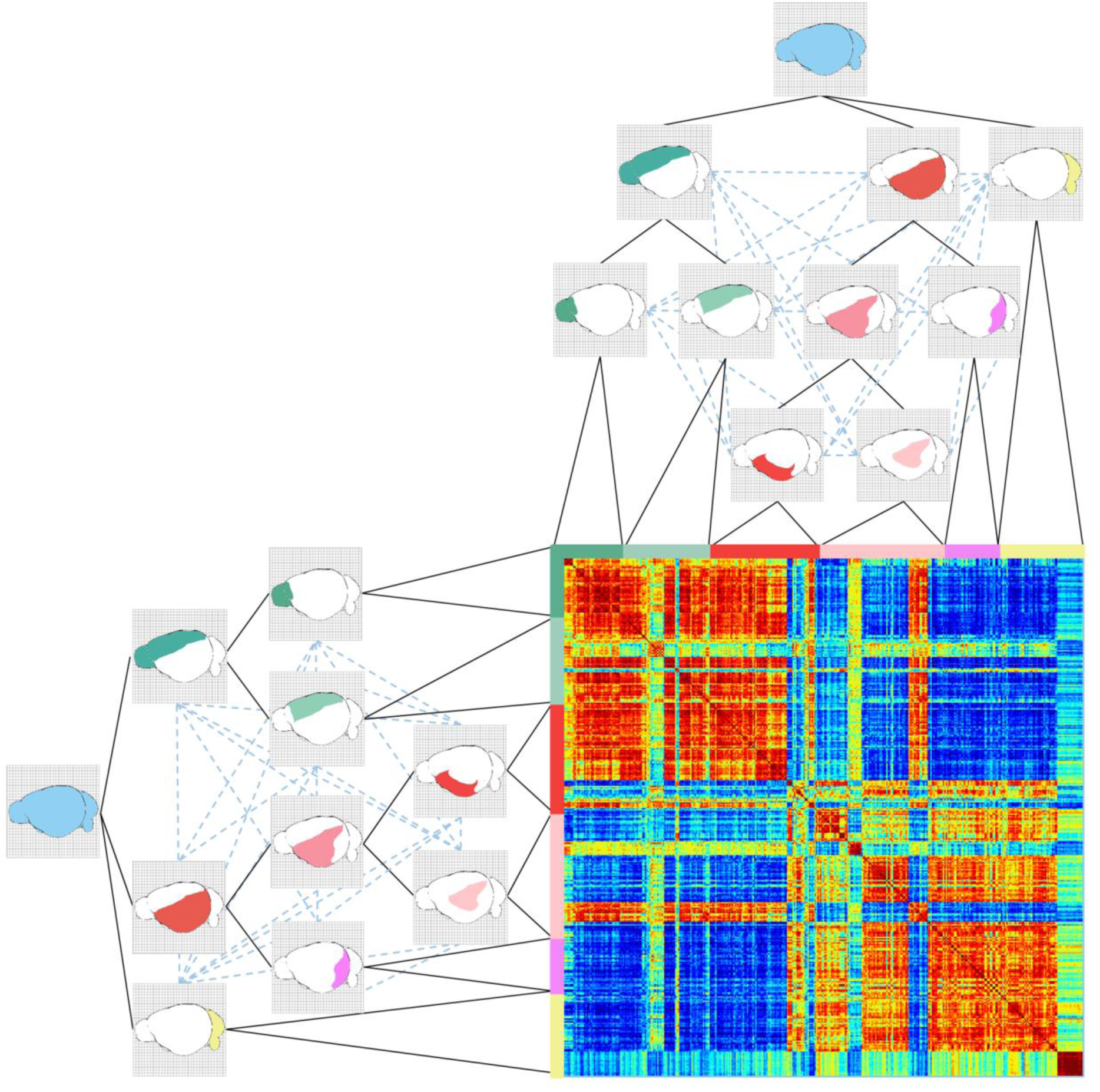
Schema of the data structure: The *Connectivity Storage* stores the connectivity on the lowest level (voxel-wise connectivity). Region-wise connectivity (dotted blue lines) is aggregated from the *Connectivity Storage* hierarchically.

#### Implementation

As central access point for the data, we created a REST API in GO (golang). It provides calls for importing data, creating caches as wells as *Aggregation Queries*. These are executed on the *Connectivity Storage*, which was implemented in C++ for memory and performance optimization. Connections are stored in a 4 byte floating point format, which supports a range of values ±1.18 × 10^−38^ to ±3.4 × 10^38^, with single precision (about 7 decimal digits). We choose this as trade-off to storage space, since higher precision would also cause higher reading times. *ROW* and *COL* indices have a 4 byte unsigned integer format, therefore the maximum amount of edges is limited by 4294967295 × 4294967295 (= 1.84467 × 10^19^). The *FILE* index associates rows with 8 byte unsigned integers to file positions, limiting the file size similarly to 1.84467 × 10^19^ connections or 64 petabyte.

We implemented two types of *Connectivity Caches*, for performance increase: A region-cache, that stores the aggregated voxel-level connectivity of lowest-level of the hierarchical brain-region parcellation, and factor *h* low-resolution versions (*h* ∈ ℕ, *h* ≤ |**I**|) of the *Connectivity Storage*, which cumulates the connectivity of *h* voxels along the Hilbert curve (basically every *h* rows of the *Connectivity Storage* are aggregated).

When executing an *Aggregation Query* for a *VOI*, ***V*** ⊆ **B**, the *Connectivity Cache Files* will be accessed first to check if the *VOI* contains cached regions **R**^**ROW** (**CACHE**)^ defined in the *ROW* index of the cache. The connectivity of a region **R** ∈ **R**^**ROW** (**CACHE**)^ will be added to the results from the cache, if **R** ⊆ **V**, i.e. all spatial positions of a region are contained within the *VOI*. Before the *Connectivity Storage* will be accessed, all *Connectivity Cache Files* will be queried until no further region in the cache can be found. Only after this, the remaining brain space positions of the VOI will be queried from the *Connectivity Storage*, so the total number of row-reads is minimized.

The anatomical hierarchy is represented in *OrientDB* (Garulli 2010), a graph database that can be used to store further region information, such as masks, 3D models or links to online repositories. Region-wise connectivities within those hierarchies, consisting of 1000-2000 regions with a maximum of 4 million edges. When querying such comparatively small graphs, the performance differences of standard graph databases to the *Connectivity Storage* is neglectable, therefore we store them in *OrientDB*, where it is directly linked to the brain regions.

To access the API, we created a web-component that allows visual queries that are based on selections of *VOI* directly in 2D slice views, visualized simultaneous in a 3D volume rendering. Via a spherical brush tool, an user-defined area can be marked. Figure 6 A shows for example a gene-expression volume, where the spherical area is drawn on voxel with high gene-expression. After selection, *Aggregation Queries* can be used to link connectivity data with volume data. The selected area (Figure 6 B), is used as input for a *Aggregation Query* on the API. The API retrieves the connectivity from the *Connectivity Storage* to all voxels that are either *targets* or *sources* from the selection, and the web component will render the connectivity as volume instantly. This represents the cumulative connectivity to (target) or from (source) the selected area (Figure 6 C). Further, the connectivity can be quantified in *Connectivity Profiles*, which shows the cumulated connectivity of the *VOI* to preselected (brain Figure 6 D).

**Figure 6.**
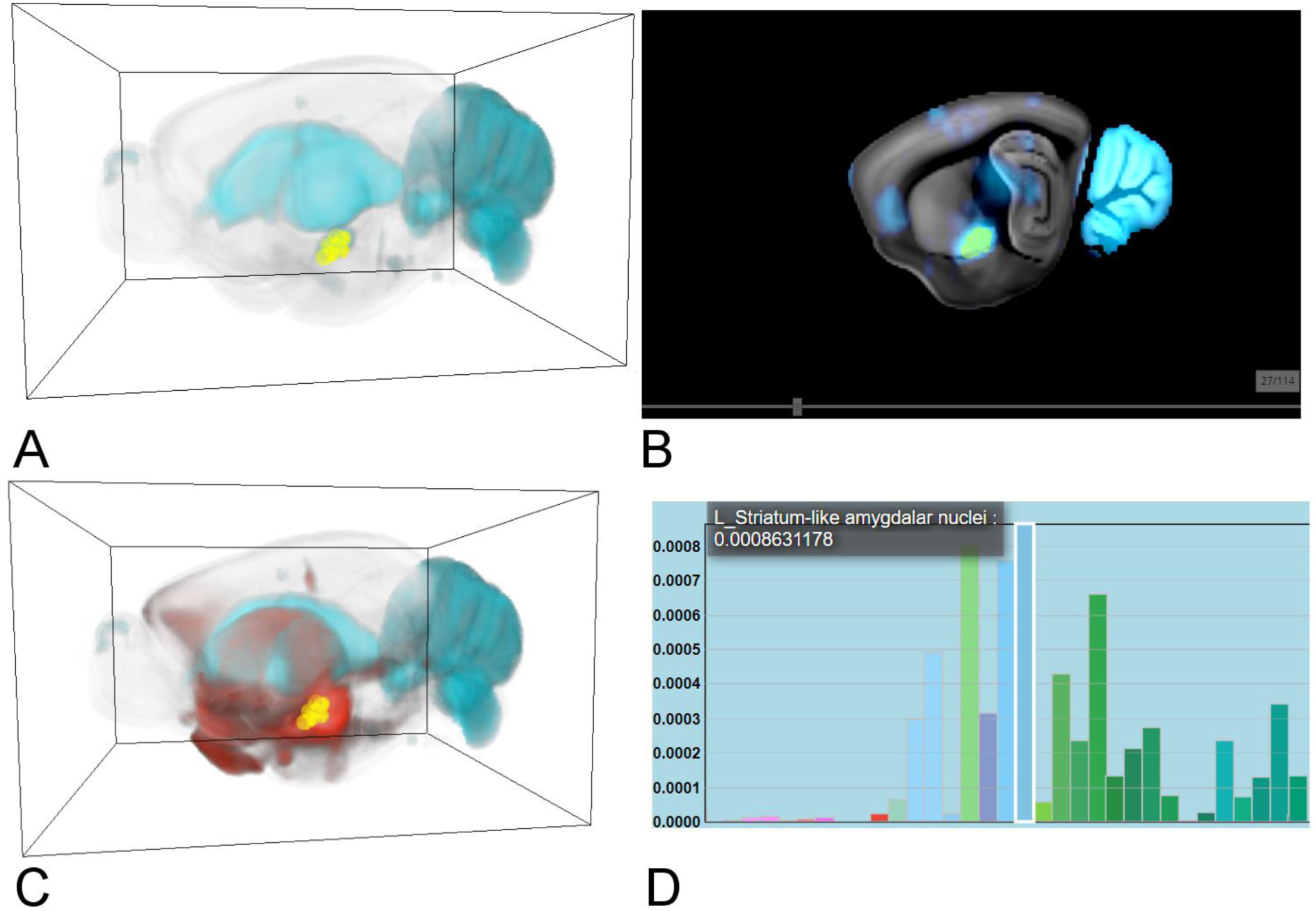
A: gene expression (cyan) with brush selection (yellow) of *VOI* in 3D ***B***: Selection has been performed on 2D slice views, ***C***: Accumulated target connectivity (red) of *VOI* in 3D ***D*:** *Connectivity Profile* of the target query, showing the mean connectivity to each brain region.

## Results

To assess the efficiency and effectivity of the data structure in context of its practical application, we performed a quantitative and qualitative evaluation on real world data introduced in *Section Data*. We quantified the effect of the data structure’s parameters (row-compression, spatial ordering, caches) on query performance and compared these results with two state-of-the-art graph engines (*Section Performance Evaluation*). We further performed two Case-Studies designed with domain experts to show relevance of the data structure for neuroscientific research (*Section Case Study 1 and 2*).

### Performance Evaluation

To verify the data structure’s applicability for real-time *Aggregation Queries*, we created test queries on three voxel-level connectivities introduced in *Section Data.* We used one directed structural connectivity matrix *SC*, resulting in two *Connectivity Storage Files* for targets and source queries, and two undirected spatial gene expression correlation networks *CS1* and *CS2* (which are further used in Case Study 1 and 2), creating one *Connectivity Storage File* each (because they are undirected).

In cooperation with domain experts, we defined 10 queries with our web-component (user-queries), which can be seen Supplementary Note 2. The *VOI* of these queries range from 0.2% to 10% of the mouse brain space. In addition, we selected 10 distinct anatomical brain regions to act as *VOI* (region-queries), their sizes ranging from 0.2% to 4% (Supplementary Note 3). To evaluate queries on a bigger scale, we further created 100 random queries, by using randomly placed spheres with random radii as *VOI*s. The sizes of these range from 0.2% to 5%, since it was not possible to place larger spheres within the mouse brain space.

We used these queries to assess the effects of individual components of the *Connectivity Storage*, such as row-compression, the spatial-ordering of rows/columns and *Connectivity Caches.* To demonstrate the data structure’s relevance for performing *Aggregation* Queries, we compared the results to the state-of-the art tools FlashGraph (Zheng et al. 2015) and GraphChi (Kyrola, Blelloch, and Guestrin 2012). We did not evaluate the performance of the *Region-wise Connectivity Database* in the *OrientDB* specifically, since retrieving a connection between two regions involves accessing only a single database entry (<10ms), in comparison to aggregating mega- to gigabytes of data from the *Connectivity Storage*.

Performance has been evaluated on an Ubuntu 16.10 64 bit machine with Intel Core i7-4470 CPU, 32 GB RAM and a 1 Terabyte SSD with a sequential read-speed of 520 MB/sec. Test on an HDD with 120 MB/sec sequential reed-speed can be found in Supplementary Note 4.

#### Effect of compressed row-storage on data size

For 3 connectivity matrices (*SC*, *CS1* and *CS2*), we created 4 *Connectivity Storage Files* (2 for *SC* and 1 for each *CS1 and CS2*). Figure 7A shows that *Connectivity Storage Files* with compression reduces the initial file size of *SC* by half, even if one is using random ordering of rows/columns. Spatial ordering by a Hilbert-curve further improves that additionally halves the sizes. The effect is smaller for *CS1* and *CS2*, since they are not as sparse as *SC* (i.e. they contain not as many zeroes).

**Figure 7.**
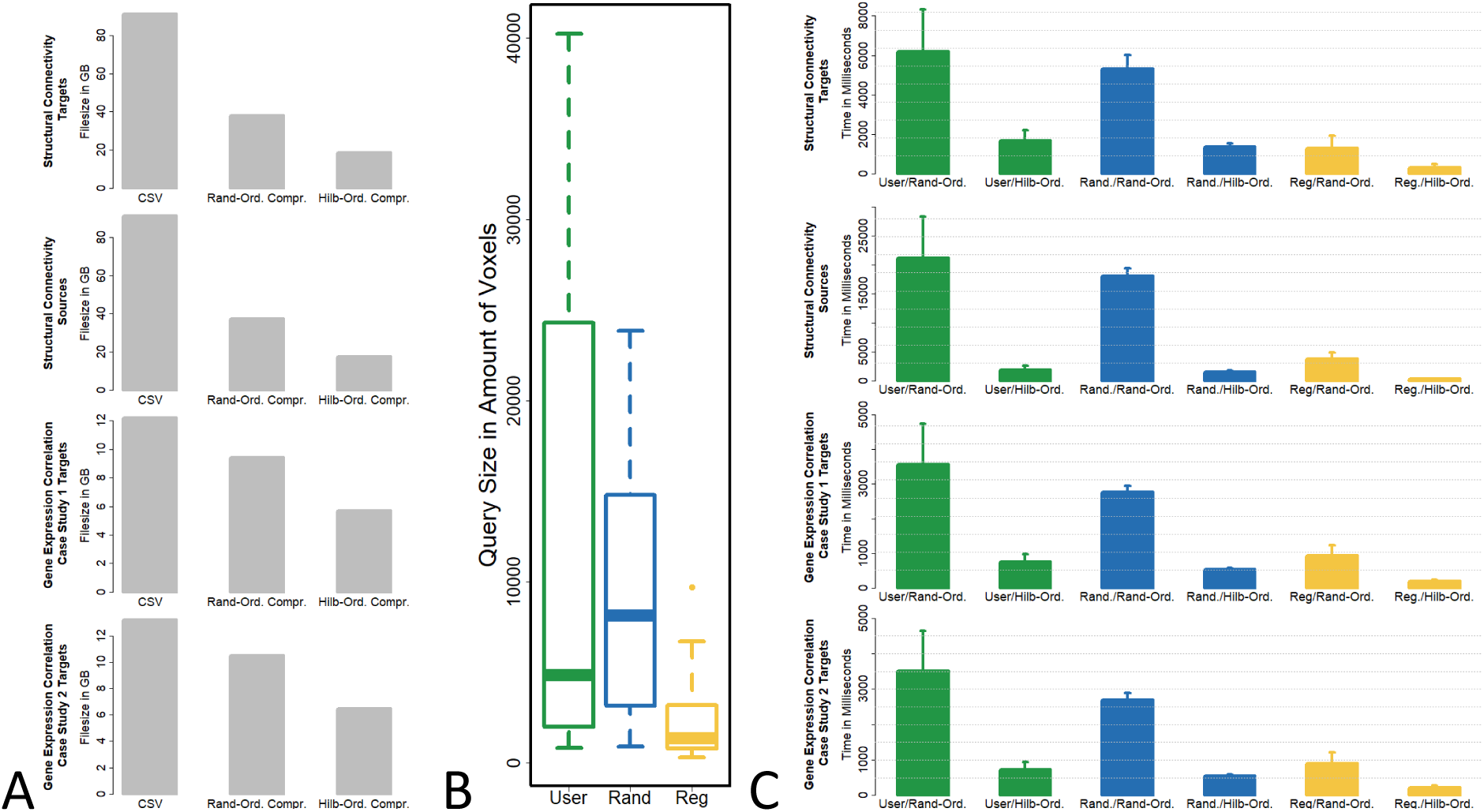
A: Effect of compressed row storage on the data size of different connectivity matrices. Bars show the size of the original CSV, the *Connectivity Storage* file without compression, random-ordering with compression and Hilbert-ordering with compression. **B:** Boxplot of the *VOI* size in amount of voxels (i.e. the query size of 10 user-defined *VOI* queries (green), 100 random *VOI* queries (blue) and 10 region *VOI* queries (yellow) **C:** Effect of spatial-ordering on query-speed on different connectivity matrices. Bars show the mean query-time with standard error of 10 user-defined *VOI* queries (green), 100 random *VOI* queries (blue) and 10 region *VOI* queries (yellow), for Hilbert-Ordering and Random-Ordering.

#### Effect of spatial-ordering on query speed

We executed the user-,random-, and region-queries on *SC*, *CS1* and *CS2* for their sources and target connectivity. Figure 7C shows the mean query time and their standard error bars on the connectivity matrices for different query types. We see, that the spatial ordering along a Hilbert curve greatly reduces query-time compared to random-ordering, especially for the bigger *SC* matrix (from up to 20 seconds to <2 seconds). This is because of read-ahead-paging, which profits from sequential reading. Note that the mean query time for different query types depends on the size of their *VOI*, so region-queries are faster than user- or random-queries simply because they involve reading fewer data (for query sizes see Figure 7B).

#### Effect of *Connectivity Caches* on query speed

As described in Section Implementation, we created a region *Connectivity Cache* of the lowest level of the hierarchical brain-region parcellation, and factor *h* low-resolution *Connectivity Caches*, where every *h* rows of the *Connectivity Storage* are aggregated, for *h* = 10 and *h* = 100. Figure 8 shows the mean query time and its standard error for different cache combinations. One can see that for high resolution *Connectivity Matrices* such as *SC*, *h*-factor caches can save up to half of the query time, while region queries profit especially from the region-caches. For lower resolutions (*CS1* and *CS2*), this effect of *h*-factor caches is not as strong. Here connectivity retrieved from *Connectivity Caches* leaves “holes” in the *VOI* of the query, so the remaining rows that need to be read from the *Connectivit*y *Storage File* are fragmented. This reduces the overall read speed, since it relies on read-ahead paging (the effect of reading spatially close rows sequentially has been shown Figure 7C). Figure 9 shows this in further detail: On the left column, one can see that the query time depends on the query size, and query time profits increasingly from *Connectivity Caches* for larger query sizes (i.e. more data to read leads to higher chances of read-ahead paging). The right column shows read-speed on *Connectivity Storage Files* vs query size and therefore the effect of reads from the *Connectivity Caches* and the resulting fragmentation. The lower read speed after cache reads is a direct cause of the higher fragmentation rate (i.e. many reads from 10-factor caches lead to more “holes” in the *VOI* than a few reads from 100-factor caches).

**Figure 8:**
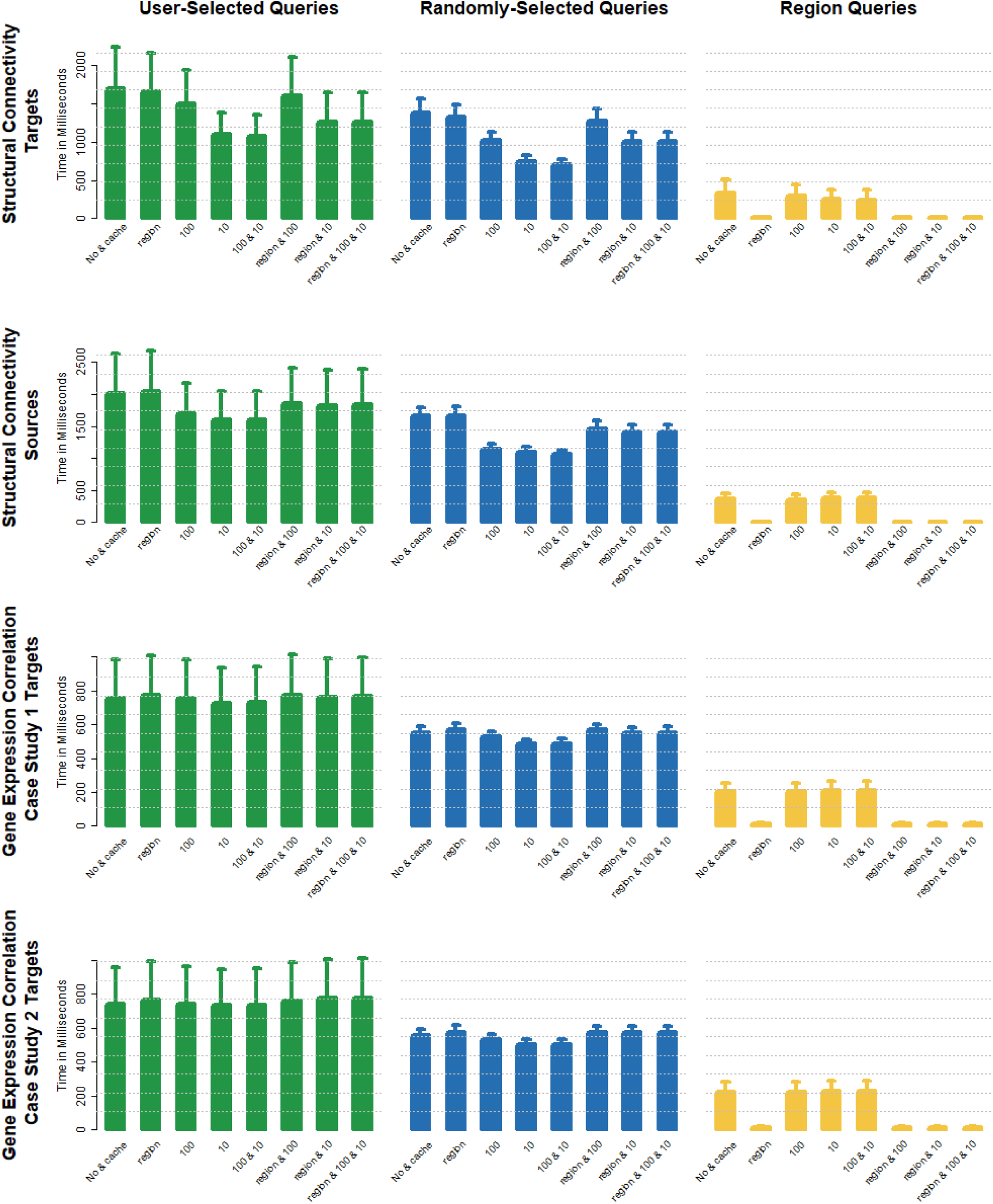
Effect of *Connectivity Cache* on query-speed on different connectivity matrices. Bars show the mean query-time with standard error of 10 user-defined VOI queries (green), 100 random *VOI* queries (blue) and 10 region *VOI* queries (yellow), for different types of caches and their combination.

**Figure 9:**
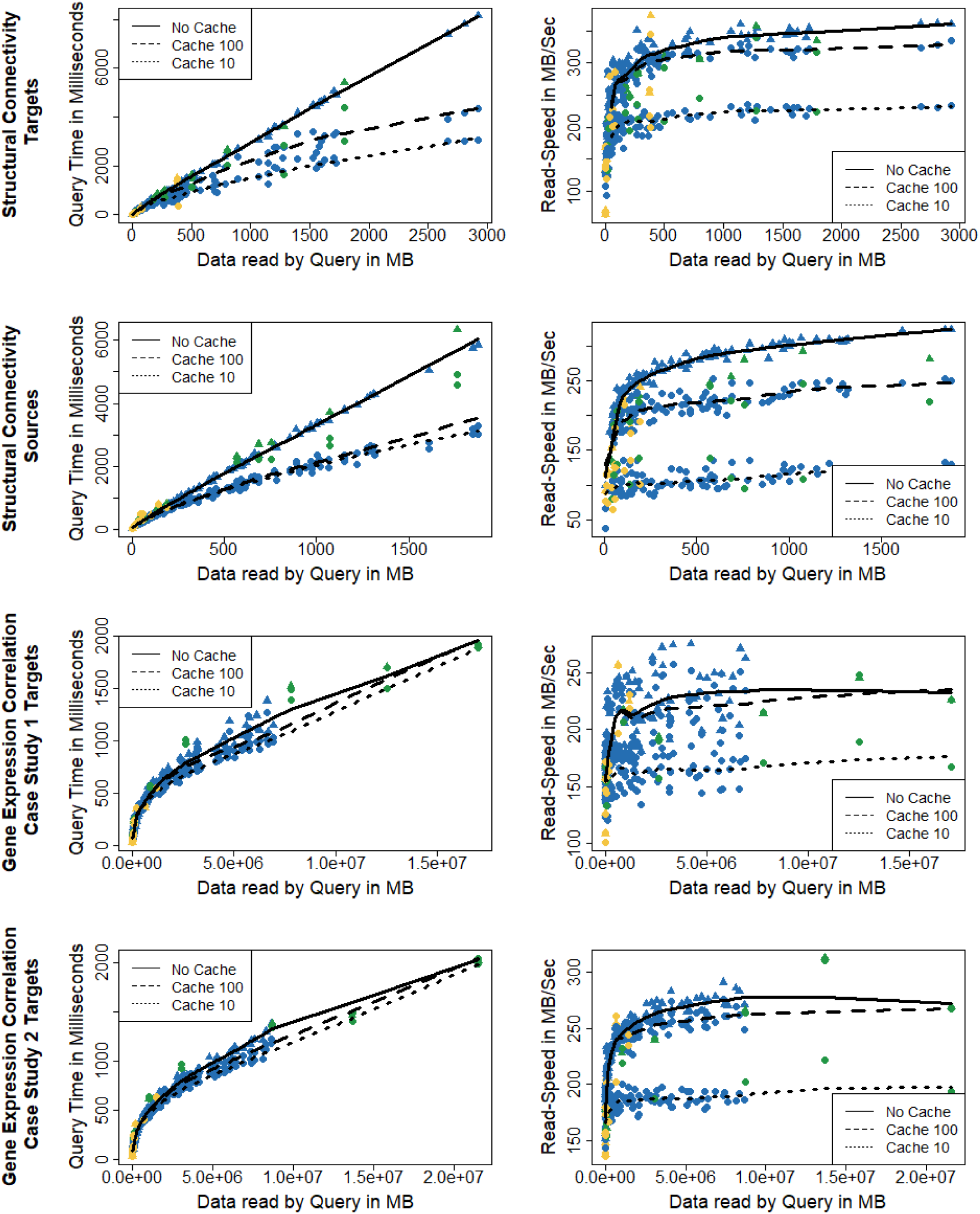
Relation of query-time and read-speed on query size for different *Connectivity Matrices, Connectivity Cache*s and query types. The left column shows the query time vs query size for queries executed with cache (**•**) and without (▲), while the color depicting the query type (green=user query, blue=random query and yellow=region query). The right column shows read speed on the *Connectivity Storage Files* vs query size, similarly encoded. LOWESS regression lines are added to see the overall trend for different cache sizes.

#### Comparison to state-of-the-art tools

We compared our method to state-of-the-art graph engines FlashGraph (Zheng et al. 2015) and GraphChi (Kyrola, Blelloch, and Guestrin 2012). Both tools are capable of computing graph algorithms (page-rank, breath-first-search etc.) on graphs with billions of edges on consumer level machines (i.e. without hundreds of gigabytes RAM). They achieve this by utilizing data access mechanisms that are able to load data from hard-drive on demand, instead of holding the whole graph in memory. GraphChi’s approach is splitting the data into small parts (so called shards), and load them on demand, while FlashGraph uses optimized IO requests for SSDs. Therefore, these methods profit at graph queries that do not involve whole graphs respectively do need to load entire connectivity matrices, such as *Aggregation Queries*. To compare their performance to the *Connectivity Storage*, we have implemented *Aggregation Queries* for both (see Supplementary Note 5 for details) and created edge-lists (<source node, target node, value>) of our connectivity matrices, which is their common input data format. Further we have ordered the node indices spatially (according to a Hilbert curve) to test them under equal conditions. Figure 10 shows that even with Hilbert ordering, FlashGraph and GraphChi do not perform as fast as our method. While on smaller graphs (CS1 and CS2), the *Connectivity Storage* is still by a factor of 2-3 faster than FlashGraph, this effect is even stronger for larger matrices (SC1), with a factor of 6. On overall, our method performs over 5x faster than FlashGraph, and 160 times faster than GraphChi. One has to note, that these tools where developed for performing various graph analysis methods, and are probably not optimized for *Aggregation Queries.* Especially GraphChi is more suited for analyzing whole graphs, while *Aggregation Queries* require only loading of subgraphs.

**Figure 10:**
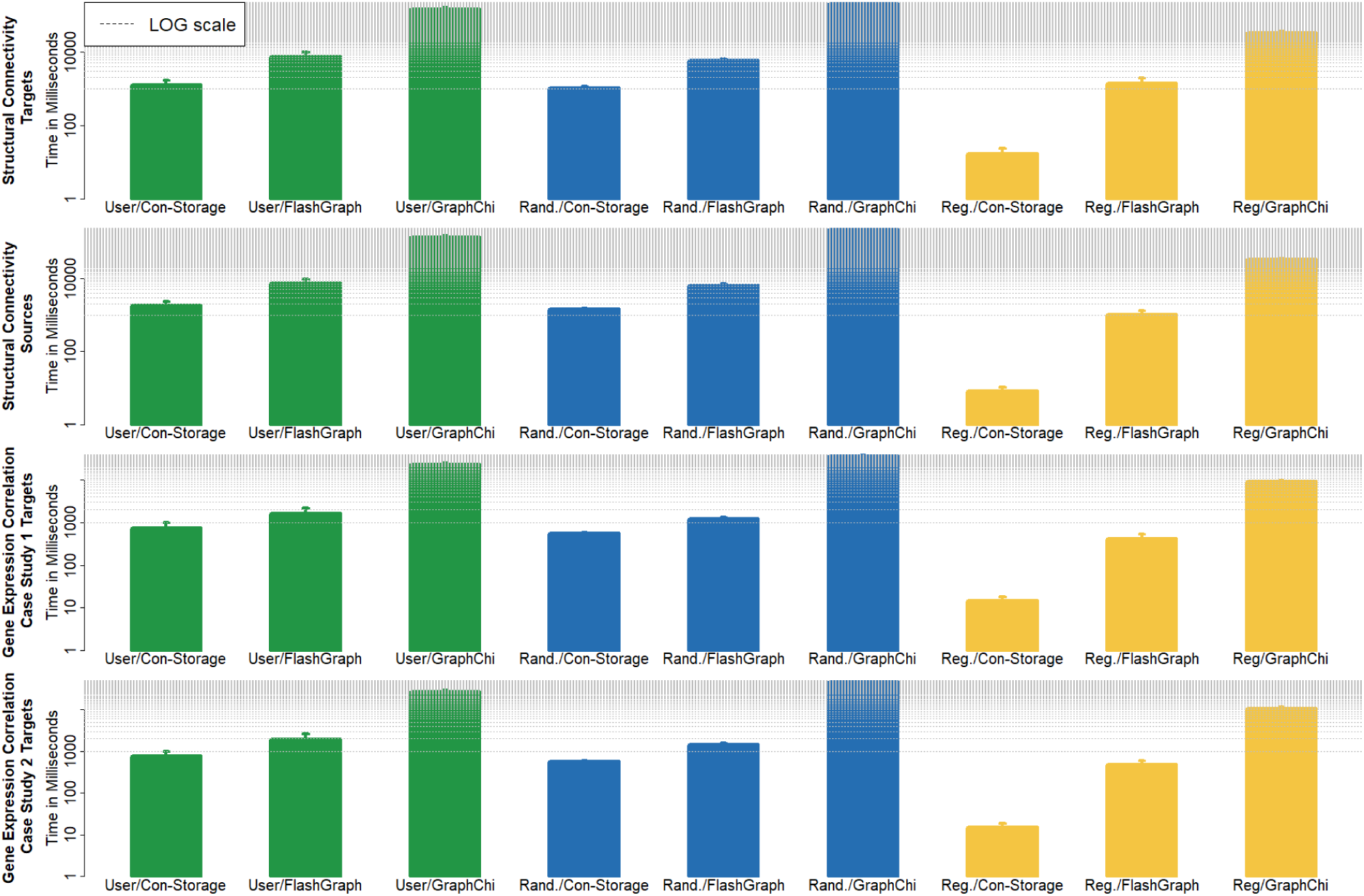
Comparison of query speed with state-of-the-art tools. Bars show the mean query-time with standard error of 10 user-defined VOI queries (green), 100 random *VOI* queries (blue) and 10 region *VOI* queries (yellow), for the Connectivity Storage, FlashGraph and GraphChi. The bars are log scaled, indicated by equidistant grey dotted lines (distance between two lines represent 1 second).

#### Example video for real-time performance

For further demonstration, Supplementary Video 1 shows a target query on the structural connectivity matrix (similar to Figure 6) performed in real-time.

### Case Study 1: Exploring different types of connectivity emerging from a brain area of interest

This case has been chosen for its particular application in circuit dissection. Recent advances in circuit neuroscience (e.g. neuro- and behavioral genetics, optogenetics, imaging) identified gene sets underlying specific behavioral function, so we mapped such function-related network context on a genetically well dissected microcircuitry (Radke 2009). To illustrate this case, we focused on the central amygdala (CEA), an amygdala subnucleus and hotspot expressing several functionally related genes, whose role in fear behavior is a heavily researched topic in the neuroscience community.

The connectivity data used for this case study consists of directed structural connectivity (*Data Set 1*) and undirected spatial gene expression correlation (*Data Set 3*), so this case demonstrates exploration of connectivities of different type and different resolution.

The entry point for our experts is a subset of these genes consisting of Prkcd (EntrezID: 18753), Sst (20604), Crh (12918), Dyn (18610) and Penk (18619) that has been known to regulate fear responses (Haubensak et al. 2010). We examined the gene expression density of these genes in 3D and 2D slice views for areas of high co-expression (where multiple genes are expressed). Image overlap of Prkdc, Crh and Dyn revealed an enclosed area (Figure 11 A, red arrow) that is selected by using a brushing tool allowing the user to interactively mark *VOI*s on 2D slice views of the brain space. We further overlayed the outlines of CEA that the selected area is indeed this brain region (Figure 11 B, red arrow).

**Figure 11.**
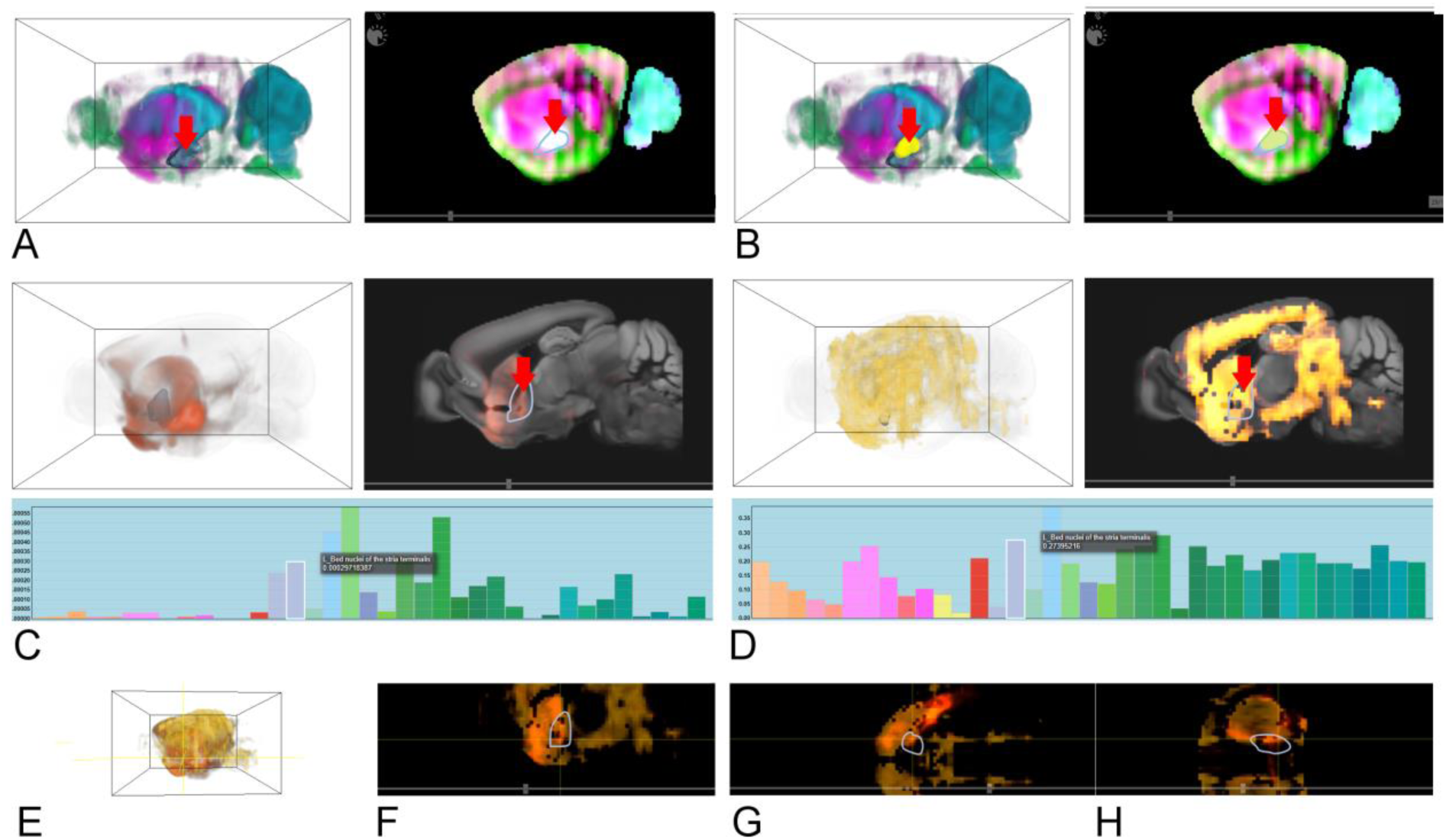
A: Overlap of Prkdc (cyan), Crh (green) and Dyn (purple) by aggregating the image intensity (i.e. strong overlap is white in the 2D slice view). Outlines of CEA in blue (red arrow). ***B***: Selecting a VOI on the image overlap (yellow), ***C:*** Structural connectivity of the VOI. Outlines of BNST in dark blue (red arrow) and its connectivity profile (bars reflect mean connectivity to (Allen Brain Atlas) brain regions, with corresponding colors). ***D***: Gene-coexpression correlation of the VOI, analogue to *C.* ***E***: Overlap of *C* and *D* in 3D. ***F***: Overlap of *C* and *D* in a 2D slice (XY). ***G***: Overlap of *C* and *D* in a 2D slice (XZ). ***H***: Overlap of *C* and *D* in a 2D slice (YZ).

After a target query on the structural connectivity (*Data Set 1*) matrix (Figure 11 C), which is performed in less than a second, particularly strong connected areas are visualized and identified by *Connectivity Profiles*. It highlights, that among other regions, the bed nucleus of the stria terminalis (BNST) has a strong connection to CEA (Figure 11, red arrow). Importantly this data confirms known structural anatomy from literature (Radke 2009). Interestingly, the BNST is functionally related to CEA. While CEA causes brief phasic fear responses, BNST shows more long-lasting tonic anxiety-like states. Thus, this approach recaptures a functional CEA-BNST circuit module for fear.

This is repeated with the spatial gene expression correlation network (*Data Set 3*) of Prkcd, Sst, Crh, Dyn and Penk, a connectivity matrix representing the voxel-wise correlation of the gene set used for this case. BNST has again the strongest connections (Figure 11 D). Figure 11 E, F, G and H visualize the overlap of both connectivity from different perspectives which demonstrates a dominant structural and genetic linkage of CEA and BNST.

### Case Study 2: Comparing networks of different modalities and species

Comparative visualization of human and animal models might be of particular interest for biomedical research and translational psychiatry. To investigate comparative functional networks across species the experts next assess this workflow by exploring functional connectivity and gene co-expression correlation from gene sets related to psychiatric traits, here exemplary autism in human (Kennedy and Adolphs 2012).

For this case study we used voxel-level undirected spatial gene expression correlation (*Data Set 3*), and region-level functional connectivity (*Data Set 2*). The data is retrieved from the *Region-Wise Connectivity Database*, which highlights the usability of our data structure on different levels of hierarchical brain parcellations. An example how the user navigates these hierarchies can be seen in *Supplementary Video 2*.

To explore and compare global gene expression correlation networks and functional MRI networks in the mouse framework, we visualized brain regions in mouse and human that are corresponding to each other (colors are picked from the *AMBA* and *AHBA* and do not correspond to each other). High coupling can be found mostly in cortex (agranular insular and temporal association areas) and primary sensory areas (olfactory, gustatory and somatosensory areas) to amygdala (central and medial).

Closer inspection of a subnetwork related to social behavior in autism, consisting of higher association cortex, namely insula cortex (IC), frontal pole (FP), hypothalamus (HY) and midbrain (MB), as well as the CEA, revealed gene co-expression correlation within the autism gene set was strongest within cortical regions (FP,IC) and weaker between cortex and subcortical structures (CEA,HY,MB). Figure 12 A shows the overlap (product) of functional connectivity and gene expression correlation of this subnetwork for mouse, Figure 12 B is analogue for human.

**Figure 12.**
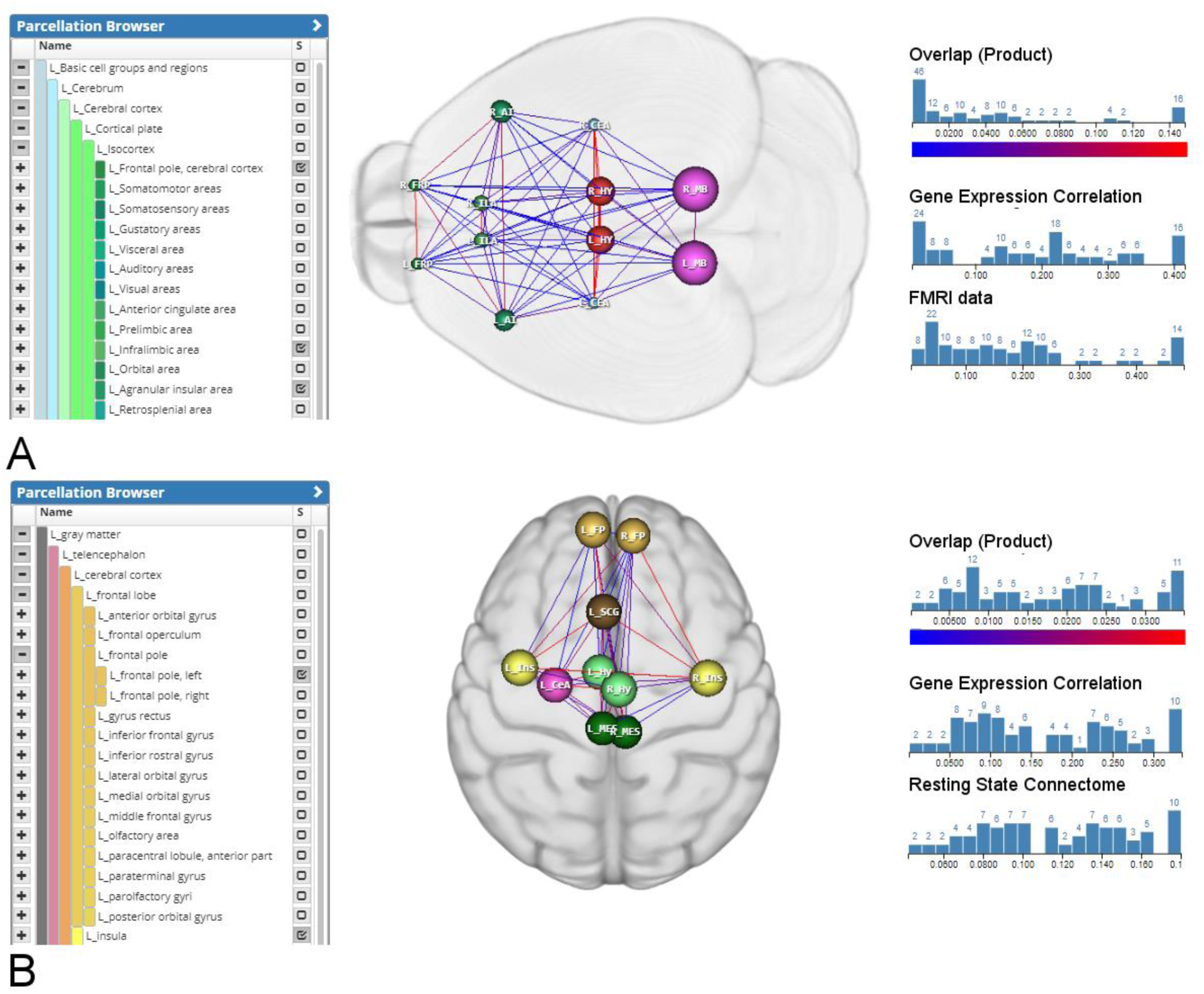
A: 2D mouse brain regions (green: cortical regions, red: HY, pink: MB and blue: CEA) and with overlap (product) of functional connectivity and gene expression correlation (blue: weak, red: strong) ***B***: 2D human brain regions (orange, brown and yellow: cortical regions, green: HY, dark-green: MB and pink: CEA), also with overlap (product of functional connectivity and gene expression correlation.

## Discussion

We have shown that our spatial connectivity data structure outperforms state-of-the-art graph engines when querying connectivity of local brain areas. To achieve even real-time access of outgoing/incoming connections without holding the whole connectivity matrices in the memory, a combination of data compression, spatial locality, memory mapping and hierarchical anatomical annotations is used.

Aggregation of outgoing/incoming connections of a brain area requires the reading of all edges of the involved network nodes. Therefore, row-wise data compression is used due the specificity of the task: It reduces the total amount of data that needs to be read from the hard drive for whole rows, but it is neglectable that it is not optimized for reading single connections. As shown in *Evaluation*, spatial organization and sparsity of the data increases compression factor by 2, compared to random ordering. Sparsity is often given in neurobiological connectomic data (Sporns 2016), and could be further improved by extended preprocessing of the data (Xu et al. 2015) in possible future projects. One has to note that row-wise compression improves only the reading speed of rows, and therefore outgoing connections. This is not an issue for undirected connectivity graphs, since those are symmetric (outgoing connections are equal to incoming connections), but requires a separate transposed *Connectivity Storage* for retrieving incoming connections. While not influencing query speed, twice the disk space is needed.

A Hilbert curve is used to generate spatial locality of rows, i.e. rows, whose nodes are spatially close in brain space are also close in the *Connectivity Storage* file. In combination with memory mapping, read-ahead paging, this greatly increases read speed, as shown in *Evaluation*. Further, it is not necessary to hold a *Connectivity Storage* file in the memory. Therefore, one can access large matrices, with billions of edges, and execute *Aggregation Queries* on multiple matrices sequentially without loading them into memory.

Depending on data acquisition techniques, neurobiological data is available on diverse scales (Betzel and Bassett 2017). To operate on different region wise levels, we used hierarchical anatomical annotations (Lein et al. 2007) and aggregated connectivity from bottom (voxel) to top (large brain regions). Since those annotations consists of only 1288, the additional stored connections are neglectable. To bridge the gap between region and voxel levels, we created a row and column indices. These allow retrieving voxel-wise data for brain regions and to map lower resolution data to a common reference space enabling comparison of connectivity of different resolution. One has to note, that this represents only upsampling of the data. Since this is done on run-time, it allows a continuous experience in visual analytic workflows, since data does not need to be preprocessed. Further, this technique can be used to create region-wise caches (voxel-wise outgoing/incoming connectivity of brain regions), or pyramids representations with lower resolution (voxel-wise outgoing/incoming connectivity of lower-resolution super voxels). Although these create additional storage overhead, we show in *Evaluation* that these greatly improve scalability, so future projects could work with even larger matrices in tera- or petabyte range.

## Conclusion

In this paper, we present a novel data structure to explore heterogeneous neurobiological connectivity data of different types, modalities and scale for interactive visual analytics workflows. It enables domain experts to combine data from large-scale brain initiatives with user-generated data, by utilizing the hierarchical and spatial organization of the data. Connectivity data at different resolutions, such as mesoscale structural connectivity and region-wise functional connectivity can be queried on different levels on a common hierarchical reference space. On the lowest level, voxel-wise brain networks with billions of edges can be accessed/queried in real-time without having them loaded into working memory. It outperforms state-of-the-art graph in receiving connectivity of local brain areas, which allows continuous interactive exploration workflows on consumer level machines and/or via web. We demonstrate this with the implementation of a web-component for visual queries, based on *VOI* selections in 2D slice views. Results are visualized in a 3D volume rendering together with brain anatomy. Case studies conducted with domain experts showed that we could reproduce findings of neural circuits research which are currently extensively investigated experimentally. An inter-species comparison of multimodal brain networks linked to autism showed even more versatile applications, and potential use in studying psychiatric conditions.

For the future, we are aiming to extend this prototype to create a holistic framework for interactive exploration of neurobiological data. This should not only allow to access the data, but also include the import, preprocessing as well as computing network statistics in the web.

## Acknowledgments

We want to thank Florian Schulze, Nicolas Swoboda, Markus Töpfer and Emre Tosun for creating and working on parts of the web-compontent. This work is result of a joint VRVis/IMP project supported by Grant 852936 of the Austrian FFG Funding Agency. VRVis is funded by BMVIT, BMWFW, Styria, SFG and Vienna Business Agency in the scope of COMET - Competence Centers for Excellent Technologies (854174) which is managed by FFG. Wulf Haubensak was supported by a grant from the European Community’s Seventh Framework Programme (FP/2007-2013) / ERC grant agreement no. 311701, the Research Institute of Molecular Pathology (IMP), Boehringer Ingelheim and the Austrian Research Promotion Agency (FFG).

## Footnotes

1 Such a multi resolution reference brain space is e.g. available from the Allen Institute, providing different kinds of data at 100-micron and 200-micron resolution.

